# Performance-optimized partitioning of clonotypes from high-throughput immunoglobulin repertoire sequencing data

**DOI:** 10.1101/175315

**Authors:** Nima Nouri, Steven H. Kleinstein

## Abstract

**Motivation:** During adaptive immune responses, activated B cells expand and undergo somatic hypermutation of their immunoglobulin (Ig) receptor, forming a clone of diversified cells that can be related back to a common ancestor. Identification of B cell clonotypes from high-throughput Adaptive Immune Receptor Repertoire sequencing (AIRR-seq) data relies on computational analysis. Recently, we proposed an automate method to partition sequences into clonal groups based on single-linkage clustering of the Ig receptor junction region with length-normalized hamming distance metric. This method could identify clonally-related sequences with high confidence on several benchmark experimental and simulated data sets. However, this approach was computationally expensive, and unable to provide estimates of accuracy for new data. Here, a new method is presented that address this computational bottleneck and also provides a study-specific estimation of performance, including sensitivity and specificity. The method uses a finite mixture modeling fitting procedure for learning the parameters of two univariate curves which fit the bimodal distributions of the distance vector between pairs of sequences. These distribution are used to estimate the performance of different threshold choices for partitioning sequences into clonotypes. These performance estimates are validated using simulated and experimental datasets. With this method, clonotypes can be identified from AIRR-seq data with sensitivity and specificity profiles that are user-defined based on the overall goals of the study.

**Availability:** Source code is freely available at the Immcantation Portal: www.immcantation.com under the CC BY-SA 4.0 license.

## 1 Introduction

Next-generation sequencing technologies are increasingly being applied to carry out detailed profiling of B cell receptors (BCRs). Identification of B cell clonotypes (sequences that are related through descent from a single naive B cell) from these high-throughput Adaptive Immune Receptor Repertoire sequencing (AIRR-seq) data relies on computational analysis. Accurate identification of clonal members is important, as these clonal groups form the basis for a wide range of repertoire analysis, including diversity analysis, lineage reconstruction, and detection of antigen-specific sequences (Yaari and Kleinstein, 2015).

Hierarchical-based clustering is a widely-used approach for partitioning sequences into clonotypes. The application of this approach has been discussed in detail by (Yaari and Kleinstein, 2015) and associated software tools have been developed by (Gupta *et al.*, 2015, 2017). Identifying clonally-related BCRs is typically accomplished in two steps. First, sequences are split into groups that share the same V-gene annotation, J-gene annotation, and number of nucleotides in their junction. Next, these groups are hierarchically-clustered based on the similarity of their junction sequence, and partitioned by cutting the dendrogram at a fixed distance threshold. We previously developed an automated approach for determining this threshold, and demonstrated that using this threshold with single-linkage clustering based on the length-normalized hamming distance detects clonotypes with high confidence on several benchmark datasets (Gupta *et al.*, 2017). However, the actual sensitivity and specificity may differ on any particular dataset, and existing methods do not provide a mechanism to estimate or tune study-specific performance.

Here, we propose and validate a computationally-efficient clustering-based method for partitioning BCR sequences into clonotypes, which allows for study-specific performance estimation. This method has been integrated into the Immcantation framework for AIRR-seq analysis, as part of the **SHazaM** R package.

## 2 Method

The proposed method extends the approach proposed by (Gupta *et al.*, 2017). Specifically, we develop a model-based method for determining the distance threshold for partitioning sequences, which allows for estimation of sensitivity and specificity. First, the “distance-to-nearest” distribution is determined using length-normalized nucleotide Hamming distance (i.e., the distribution of minimum distances from each sequence to every other non-identical sequence). This is typically a bimodal distribution, with the first mode representing sequences with clonal relatives and the second mode representing those without clonal relatives (i.e., singletons) in the dataset (Figure 1 panel a.2). Next, this distribution is explicitly modeled as a mixture of two univariate distribution functions (e.g. a mixture of Gamma or Normal distribution) of the form:

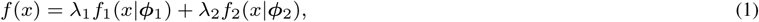

where *λ* _1_ and *λ* _2_ represent the mixing weights (sum to one), *x* represents the nearest neighbor distance, and *φ* represents the vector of each component parameters. Here, we investigate all combinations of *f*_1_ and *f*_2_ as Gaussian and Gamma distributions so *°* is either the mean and standard deviation (*μ, σ*) of a Normal distribution, or the shape and scale (*k, θ*) of a Gamma distribution. A maximum-likelihood fitting procedure (function `fitdistr` from package **MASS**) is used to estimate the parameters of the model as follows: (1) parameters of the model are initialized using a standard Gaussian mixture model (GMM). The GMM estimates mixing weight *λ*_*i*_, mean *μ*_*i*_ and standard deviation *σ*_*i*_ where *i ∊* {1, 2} refers to the first and second distributions. (2) These parameters are then used as initial values to begin the fitting procedure (if Gamma distribution is chosen, the initial values are translated accordingly).

After fitting, the two distributions are used to estimate sensitivity (SEN) and specificity (SPC) by the fractions TP/(TP+FN) and TN/(TN+FP), respectively. The statistical rates (true positive (TP), false negative (FN), false positive (FP), and true negative (TN)) are given by the area under the curves:

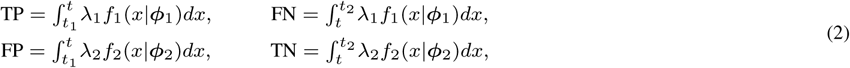

where *t*_1_ and *t*_2_ are the minimum and maximum values of the distance-to-nearest distribution, respectively.

**Fig. 1.**
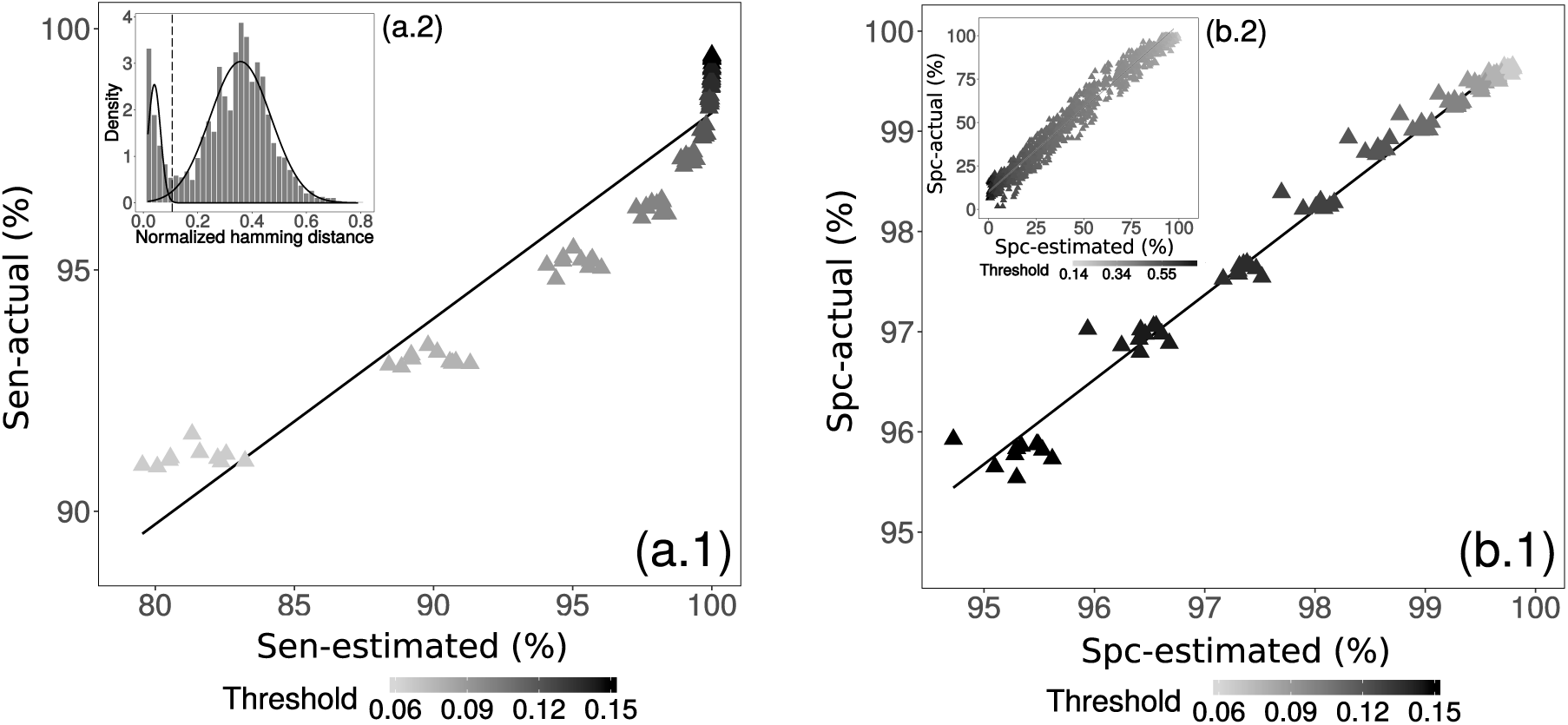
Performance assessment of (a) sensitivity and (b) specificity for determining clonal relatedness. Mixture modeling of the distance-to-nearest distribution (panel a.2) was used to estimate sensitivity and specificity for each specified value of the threshold (vertical line in a.2) according to equation set (2). The estimated performance was compared with actual performance for simulated (a.1 and b.1) and experimental (b.2) data across a wide range of thresholds (shades of grey for each point). Note that the choice of bin size impacts the shape of plotted histogram in (a.2), while the fitting procedure is independent of this bin size. Panel b.2 shows representative performance from one base individual.

## 3 Results

To determine the optimal distributions to use for the mixture model, we evaluated all four combinations of Gaussian and Gamma distributions for *f*_1_ and *f*_2_. For each combination, the log likelihood was determined for 10 simulated (set R1 generated by (Gupta *et al.*, 2017)), and 73 experimental (dengue infected individuals from (Parameswaran *et al.*, 2013)) datasets. We found that in 80% of trials the choice of Gamma distribution for both *f*_1_ and *f*_2_ yielded the highest likelihood. Further, in each trial, visual inspection suggested that this choice placed the threshold approximately equidistant between the two distributions. Therefore, Gamma distributions were selected as the default choices for *f*_1_ and *f*_2_, and used in all of the analyses below.

The ability of the proposed method to estimate sensitivity and specificity for clonal relatedness was evaluated on both simulated and experimental data. First, sensitivity and specificity were evaluated using ten simulated datasets (set R1 generated by (Gupta *et al.*, 2017)). On each dataset, a wide range of potential thresholds for partitioning sequences into clonotypes were considered. At each threshold value, we calculated the actual performance based on the known clonal relationships from the simulation, as well as the estimated performance based on the mixture modeling and equation set (2) using the area under the fitted distribution curves. We found a high correlation between the actual and estimated sensitivity (*R*^2^ = 92%) and specificity (*R*^2^ = 98%) on average over all ten simulated datasets (Figure 1, panel (a.1) and (b.1)).

The specificity of the methodology was further evaluated using experimental BCR sequencing from 73 individuals with acute dengue infection (Parameswaran *et al.*, 2013). First, one of the individuals was chosen as the “base". Next, a single sequence was chosen randomly from each of the remaining individuals and added to the sequencing data from the base individual. Specificity was then defined by how often the sequences from non-base individuals were correctly determined to be singletons. Any grouping of these sequences into larger clonotypes must be a false positive since, by definition, a B cell clone cannot span two individuals. Like the simulated data, specificity was calculated both using the known source of the sequences (actual) and for the mixture model estimate. This procedure was then repeated twenty times for each of ten different base individuals (chosen randomly). The results indicated a high correlation between the actual and estimated specificity (*R*^2^ = 95%) across all ten base individuals (Figure 1 panel b.2). Overall, the results on the simulated and experimental datasets indicate that the mixture modeling method provides an accurate estimate of sensitivity and specificity for partitioning sequences into clonotypes.

## 4 Conclusion

We have proposed and validated a computationally-efficient clustering-based method for automatically partitioning BCR sequences into clonally-related groups. The method is based on a mixture model fit to the bimodal distance-to-nearest distribution, and allows for direct estimation of the sensitivity and specificity for membership in a multi-sequence clone. The ability to estimate sensitivity and specificity directly from a BCR sequencing dataset allow researchers to identify B cell clonotypes with performance characteristics that optimize study-specific goals. For instance, a threshold with high-sensitivity may be ideal for identifying sequences that are part of a clone expansion including a known antigen-specific sequence, while a threshold with high-specificity may be ideal for determining biological connections between tissue compartments or B cell subsets. As a default, the threshold is chosen to maximize the average of sensitivity and specificity. This new procedure has been implemented under the findThreshold function as part of the SHazaM R package in the Immcantation framework: www.immcantation.com.

## Acknowledgements

The authors thank N. T. Gupta for providing the simulated dataset, and S. Marquez for useful comments on code. This work was supported by the HPC facilities operated by, and the staff of, the Yale Center for Research Computing. This work was supported in part by the National Institutes of Health (NIH) under award number R01AI104739.

